# The NHR-23-regulated putative protease inhibitor *mlt-11* gene is necessary for *C. elegans* cuticle structure and function

**DOI:** 10.1101/2024.05.12.593762

**Authors:** James Matthew Ragle, Ariela Turzo, Anton Jackson, An A. Vo, Vivian T. Pham, Jordan D. Ward

## Abstract

*C. elegans* molting offers a powerful entry point to understanding developmentally programmed apical extracellular matrix remodeling. However, the gene regulatory network controlling this process remains poorly understood. Focusing on targets of NHR-23, a key transcription factor that drives molting, we confirmed the Kunitz family protease inhibitor gene *mlt-11* as an NHR-23 target. Through reporter assays, we identified NHR-23-binding sites that are necessary and sufficient for epithelial expression. We generated a translational fusion and demonstrated that MLT-11 is localized to the cuticle and lined openings to the exterior (vulva, rectum, mouth). We created a set of strains expressing varied levels of MLT-11 by deleting endogenous *cis-*regulatory element sequences. Combined deletion of two *cis-*regulatory elements caused developmental delay, motility defects, and failure of the cuticle barrier. Inactivation of *mlt-11* by RNAi produced even more pronounced defects. *mlt-11* is necessary to pattern every layer of the adult cuticle, suggesting a broad patterning role prior to the formation of the mature cuticle. Together these studies provide an entry point into understanding how individual *cis-*regulatory elements function to coordinate expression of oscillating genes involved in molting and how MLT-11 ensures proper cuticle assembly.

## INTRODUCTION

Molting is an essential process in ecdysozoans in which a new exoskeleton is built and the old one is shed, allowing animals to grow in size. Molting has also been highlighted as a drug target to combat parasitic nematodes, a group of pathogens that infect over a billion humans and infests crops and livestock (Kumar et al. 2007; Charlton et al. 2010; Page et al. 2014). It is a developmental process unique to arthropods and nematodes and involves druggable molecules, such as nuclear hormone receptors and proteases, some of which are nematode-specific. In contrast to arthropods, nematode molting is poorly understood.

Following hatching, nematodes progress through four larval stages before reaching adulthood. During each larval stage, there is extensive secretion of the materials required to build a new apical extracellular matrix (cuticle)(Lažetić and Fay 2017; Sundaram and Pujol 2024). The cuticle is a multi-layered structure largely composed of collagens and cuticlins. The outermost layer (glycocalyx/ surface coat) is rich in carbohydrates and mucins that cover a thin layer of lipids and glycolipids (epicuticle) (Lažetić and Fay 2017). A specialized, transient structure known as the pre-cuticle is thought to pattern the new cuticle and is then endocytosed (Cohen and Sundaram 2020; Sundaram and Pujol 2024). Towards the end of each stage, animals enter a sleep-like phase called lethargus where the old cuticle begins detaching from the epidermis, termed apolysis (Singh and Sulston 1978; Lažetić and Fay 2017). Animals then undergo ecdysis, which involves a series of longitudinal contractions and expansions and forward thrusts to escape the old cuticle (Singh and Sulston 1978; Lažetić and Fay 2017).

*C. elegans* molting is controlled by a poorly understood gene regulatory network. Approximately 20% of *C. elegans* genes oscillate, peaking at distinct points in each larval stage. This oscillator is thought to coordinate aECM assembly during molting (Kim et al. 2013; Hendriks et al. 2014; Meeuse et al. 2020). A screen of 92 oscillating transcription factors identified six whose depletion altered the number or duration of molting, two of which were novel regulators (BED-3, GRH-1)(Meeuse et al. 2023). It also identified BLMP-1, which has been implicated in timely molting as a pioneer factor controlling the duration and amplitude of oscillatory gene expression (Hauser et al. 2021 Jul 5; Stec et al. 2021). Other factors previously linked to molting, include the myelin regulatory family transcription factor MYRF-1/PQN-47 and the nuclear hormone receptors, NHR-23 and NHR-25 (Kostrouchova et al. 1998; Gissendanner and Sluder 2000; Kostrouchova et al. 2001; Gissendanner et al. 2004; Frand et al. 2005; Russel et al. 2011; Meng et al. 2017; Johnson et al. 2023). However, how these factors promote oscillatory gene expression with different phases is poorly understood.

We focused on NHR-23 as it coordinates factors involved in molting, aECM remodeling, and lipid transport/metabolism (Johnson et al. 2023). It also plays a separate role promoting spermatogenesis (Ragle et al. 2020; Ragle et al. 2022 Sep 22). We were particularly interested in NHR-23 regulation of proteases and protease inhibitors as these are druggable molecules and their role in *C. elegans* aECM remodeling is not well understood. *mlt-11* is a poorly characterized gene with 10 Kunitz protease inhibitor domains and RNAi is reported to produce ecdysis defects (Frand et al. 2005; Ragle et al. 2022). *nhr-23* regulated the expression of a *mlt-11* promoter reporter and there are six NHR-23 ChIP-seq peaks flanking and within the *mlt-11* gene body (Frand et al. 2005; Gerstein et al. 2010; Johnson et al. 2023). However, the relative role of these potential *cis-*regulatory elements in controlling *mlt-11* expression was not known. Here we explore the role of the four candidate *cis-*regulatory elements in the *mlt-11* promoter using reporter assays, a *mlt-11::mNeonGreen* translational fusion, endogenous promoter deletions, and phenotypic assays.

## RESULTS

### *mlt-11* is expressed in embryonic and larval epidermal cells

We had previously used a 2.8 kb *mlt-11* promoter fragment to drive robust hypodermal expression of *nuc-1::mCherry* (Clancy et al. 2023 Feb 7). Using this 2.8 kb promoter fragment, we created a single copy *mlt-11p::NLS::mNeonGreen* promoter reporter. We observed expression in embryos starting at the bean stage in posterior epithelial cells. This persisted through the 3-fold stage spreading more anteriorly (Fig. 1A). We also detected expression in hypodermal and seam cells, similar to previous reports using an extrachromosomal array-based promoter reporter (Fig. 1B)(Frand et al. 2005). We detected expression in rectal and vulval cells in both larvae and adults (Fig. 1B). A previous promoter reporter used 3 kb of sequence upstream separated from the transcription start site (TSS) by about 2 kb of sequence (Frand et al. 2005). We created a single copy *mlt-11p::NLS::mNeonGreen* promoter reporter containing this 3 kb of sequence as well as an additional reporter containing sequence spanning from the TSS to 5.3 kb upstream (Fig 1C). Similar expression timing and intensity was seen in these two reporters compared to our original 2.8 kb promoter reporter (Fig 1C).

**Fig. 1.**
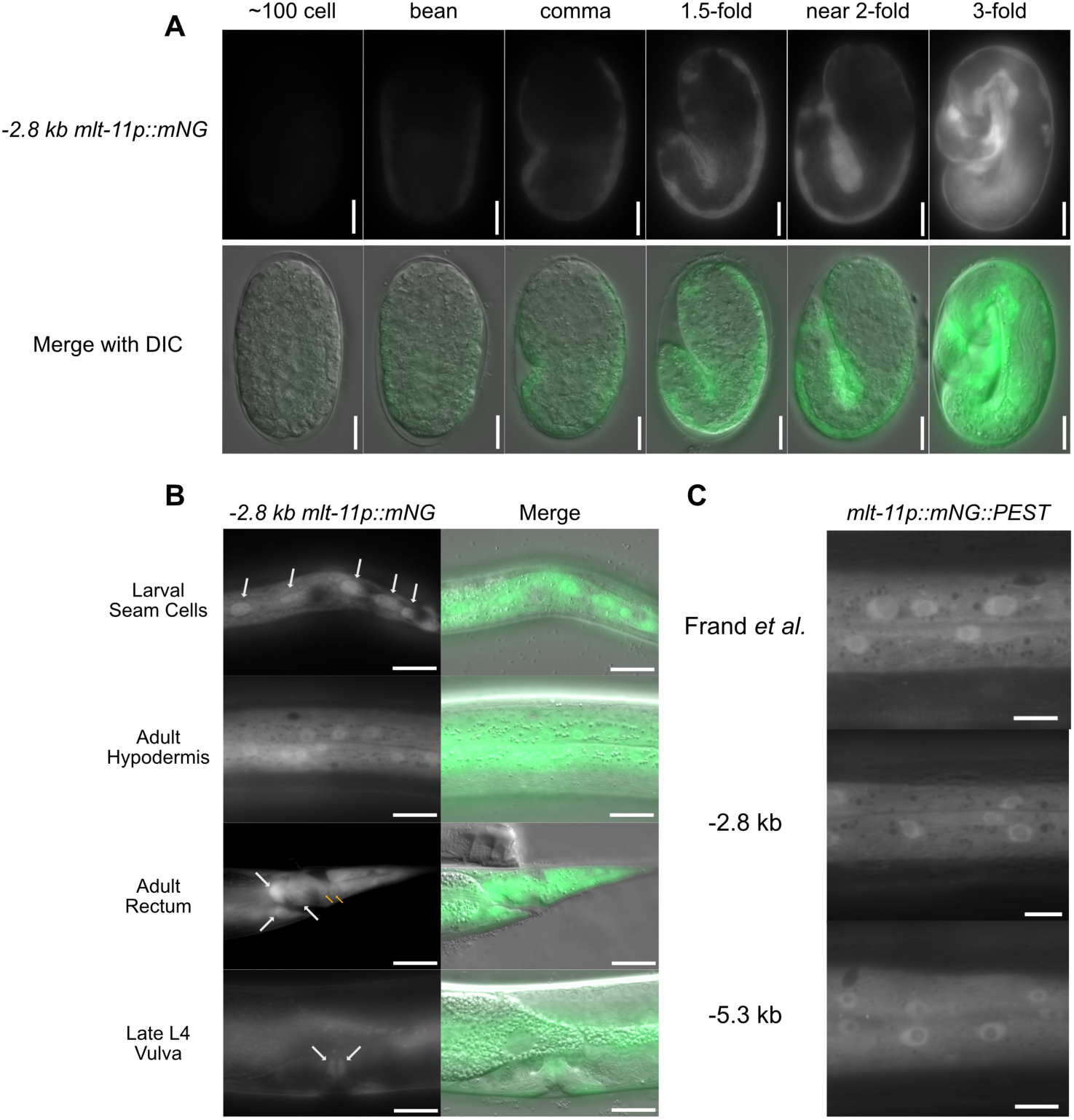
*mlt-11* is expressed in larval and embryonic epidermal cells. (A) Time course of *-2.8kb mlt-11p::mNG::tbb-2 3’UTR* expression in embryos with stages determined by embryo morphology. (B) Expression pattern of a *-2.8kb mlt-11p::mNG::tbb-2 3’UTR* promoter reporter in hypodermal cells of L4 worms. White arrows indicate seam cells (top), rectal epithelial cells (middle) and vulval cells (bottom) and yellow arrows indicate hypodermal cells near rectum. Images are representative of 40 animals examined over two biological replicates. (C) Hypodermal expression pattern in L4s of three single-copy integrated promoter reporters. One reporter uses the same sequence used by Frand *et al*. (2005) and the other two use −2.8 kb and 5.2 kb of sequence upstream of the *mlt-11* transcriptional start site (TSS).

### NHR-23 regulates *mlt-11* transcriptionally by binding multiple regions of the *mlt-11* promoter

Given the similar expression levels of the promoter reporters, we wanted to explore potential regulatory elements in the overlapping sequence of our 2.8 kb promoter reporter and the Frand *et al*. reporter. There are four NHR-23 ChIP-seq peaks in the *mlt-11* promoter (Gerstein et al. 2010; Johnson et al. 2023), and the sequences under these peaks are highly conserved in other nematodes (Fig. 2A; see Conservation track). There are also single NHR-23 peaks in the gene body and 3’ UTR, which we did not pursue further. There is a strong NHR-23 ChIP-seq peak (Peak 3) contained in both the Frand promoter reporter and our 2.8 kb promoter reporter (Fig. 2A). The Frand *et al*. reporter contained NHR-23 ChIP-seq peak 4 (Peak 4) and our reporter contained NHR-23 ChIP-seq peaks 1 and 2 (Fig. 2A). We therefore tested whether the sequence contained in each NHR-23 ChIP-seq peak, as well as two candidate enhancers identified by ATAC-seq were sufficient to drive the expression of a reporter containing a *pes-10* minimal promoter. The *pes-10Δ::mNeonGreen* transgene displayed undetectable expression in the absence of an added *cis-*regulatory element (Fig. 2B). When the sequences from peaks 3 and 4, respectively, were added we detected reporter expression in hypodermal cells with the peak 3 reporter having the most robust expression (Fig. 2B). Additionally, reporters containing the sequences from peaks 2, 3 or 4 all expressed in seam cells (Fig. 2B). We could not detect expression of reporters containing the sequences from peak 1 nor ATAC-seq peaks 1 or 2 (Fig. 2B). To test whether NHR-23 regulated expression of the *mlt-11 peak 3 pes-10::mNeonGreen* promoter reporter, we performed *nhr-23* RNAi. RNAi of *nhr-23* reduced expression of both full length (Fig. 2C) and *mlt-11 peak 3 pes-10::mNeonGreen* promoter reporters (Fig. 2C). These data indicate that NHR-23 regulates *mlt-11* and that the DNA sequences in NHR-23 ChIP-seq peaks 3 and 4 play an important role in this regulation.

**Fig. 2.**
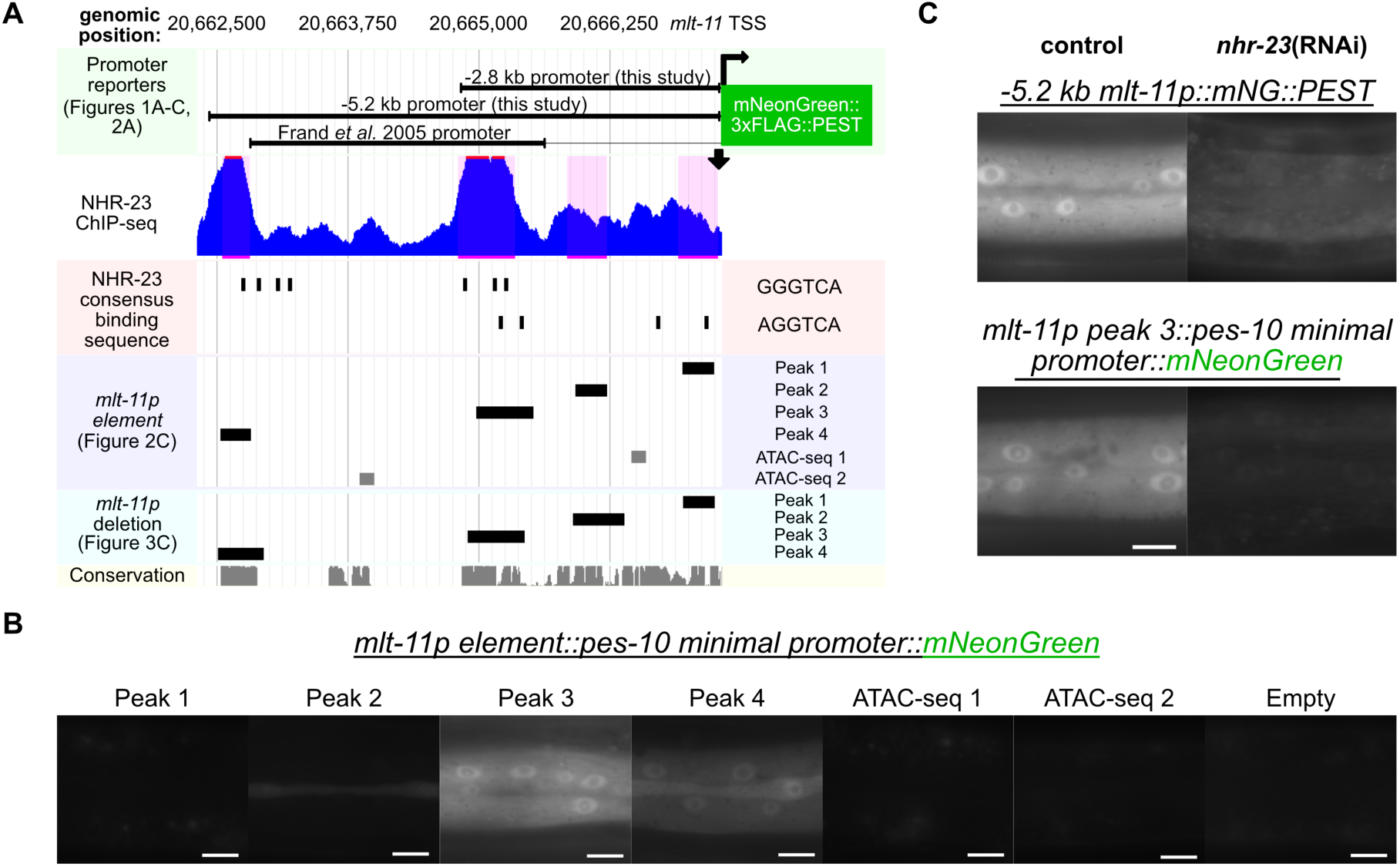
NHR-23 occupied regions in *mlt-11* promoter are sufficient to drive reporter expression. (A) Genome browser track of *mlt-11* promoter depicting NHR-23 ChIP-seq peaks; relative genomic locations of NHR-23 binding motifs; relative genomic locations of *mlt-11* promoter reporters used in this study; relative genomic locations of candidate *cis-*regulatory elements from the *mlt-11* promoter inserted into a *pes-10* minimal promoter in this study; conservation calculated across 26 nematode species. (B) Hypodermal expression pattern in L4s of single-copy integrated promoter reporters with the *pes-10* minimal promoter elements combined with *mlt-11* promoter sequence fragments corresponding to either NHR-23 ChIP-seq or peaks or ATAC-seq open chromatin regions. (C) Hypodermal expression pattern in L4 larvae of the indicated single-copy integrated promoter reporters. Reporter strains were fed either control or *nhr-23(RNAi)* knockdown conditions. Scale bars are 20 μm in B, D, E and F and 10 μm in C.

### NHR-23 ChIP-seq peaks 3 and 4 are necessary for endogenous MLT-11 expression

We next wanted to test whether the NHR-23 occupied regions from the ChIP-seq dataset were necessary for endogenous MLT-11 expression. To monitor expression we knocked in an *mNeonGreen::3xFLAG* cassette into the 3’ end of the 7th *mlt-11* exon producing an internal translational fusion that labels all described *mlt-11* isoforms (Fig. 3A). MLT-11::mNeonGreen::3xFLAG (MLT-11::mNG) was detected consistently throughout larval development in lysosomes of the hypodermis as determined by vesicle morphology and size (Miao et al. 2020) as well as in the aECM of hyp7 and seam cells (Fig. 3B). MLT-11::mNG was also observed in the epithelial aECM of other external-facing orifices: the buccal cavity, vulva, excretory duct and rectum (Fig. 3B). Using CRISPR/Cas9, we then individually deleted the sequences from each NHR-23 ChIP-seq peak in the MLT-11::mNG background and observed expression of the translational fusion (Fig. 3C). We observed the expression of MLT-11::mNG to be comparable to the control following deletion of the sequences from peaks 1 or 2, but reduced with deletion of the sequences from peaks 3 and 4. Simultaneous deletion of the sequences from peaks 3 and 4 in the same strain resulted in severely reduced expression of MLT-11::mNG, suggesting these sequences are both necessary for activation of expression of *mlt-11* by NHR-23. We also scored developmental rate, motility, and size (Fig. 4). Interestingly, *peak 2Δ*, *peak 3+4Δ* and *mlt-11(RNAi)* animals exhibited comparable developmental delay (Fig. 4A), but *peak 2Δ* animals had wild type motility and size (Fig. 4B,C). In contrast, peak 3+4Δ and *mlt-11(RNAi)* animals exhibited a significant motility defect with the RNAi causing a stronger phenotype and a comparable smaller body size (Fig. 4B,C).

**Fig. 3.**
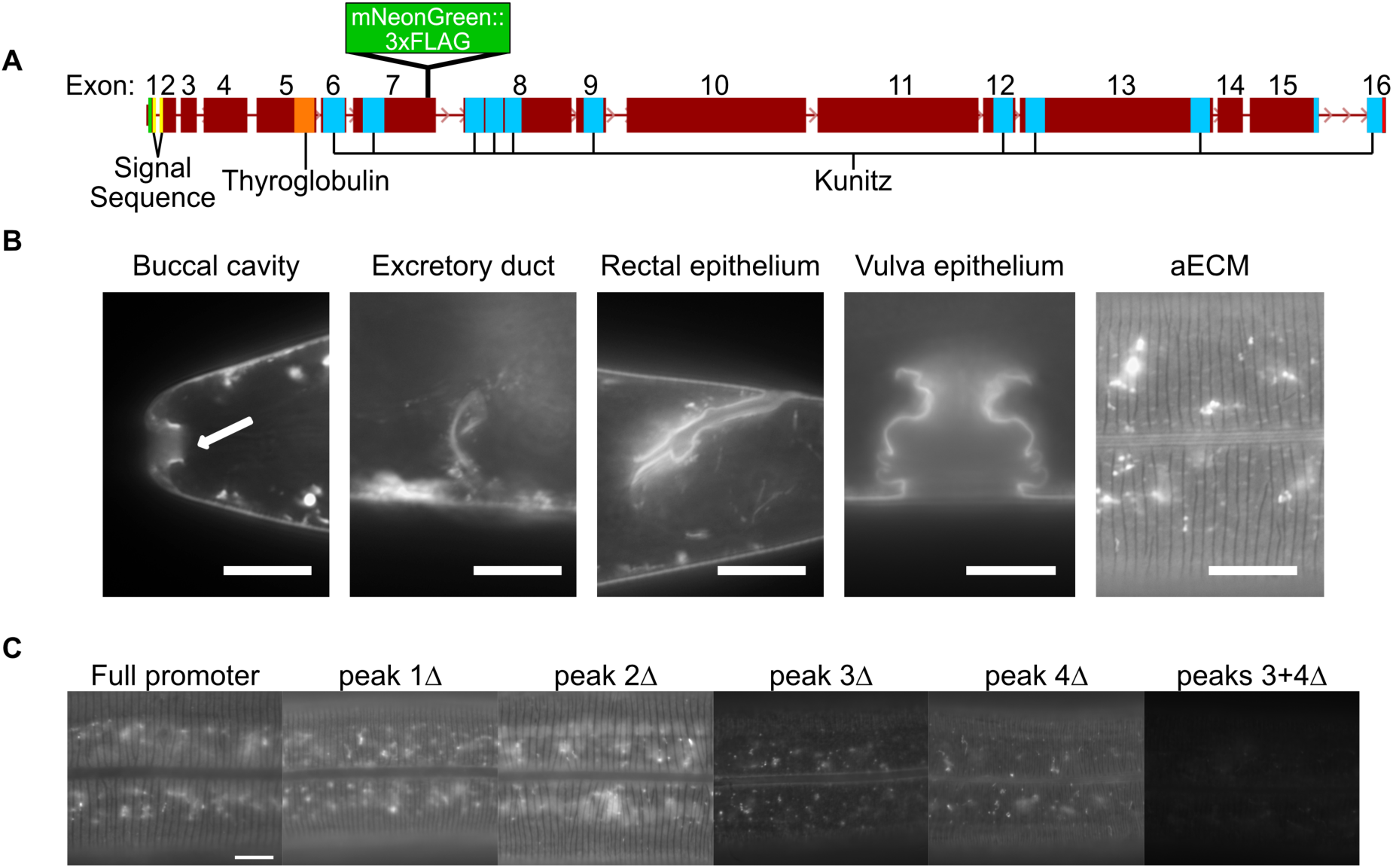
MLT-11 localizes to the aECM in seam and hypodermal cells and interfacial orifices. (A) Schematic of the internal *mlt-11::mNG::3xFLAG* knock-in. (B) Representative images of MLT-11::mNG localization in the lumen of the buccal cavity, excretory duct and rectum. MLT-11 also to the aECM and to internal vesicles in seam and hypodermal cells. Images are representative of 40 animals examined over two biological replicates at stage L4.5. Scale bars are 20 μm. (C) Representative images of MLT-11::mNG::3xFLAG in mid-stage L4 larvae with endogenous deletion of the indicated promoter regions

**Fig. 4.**
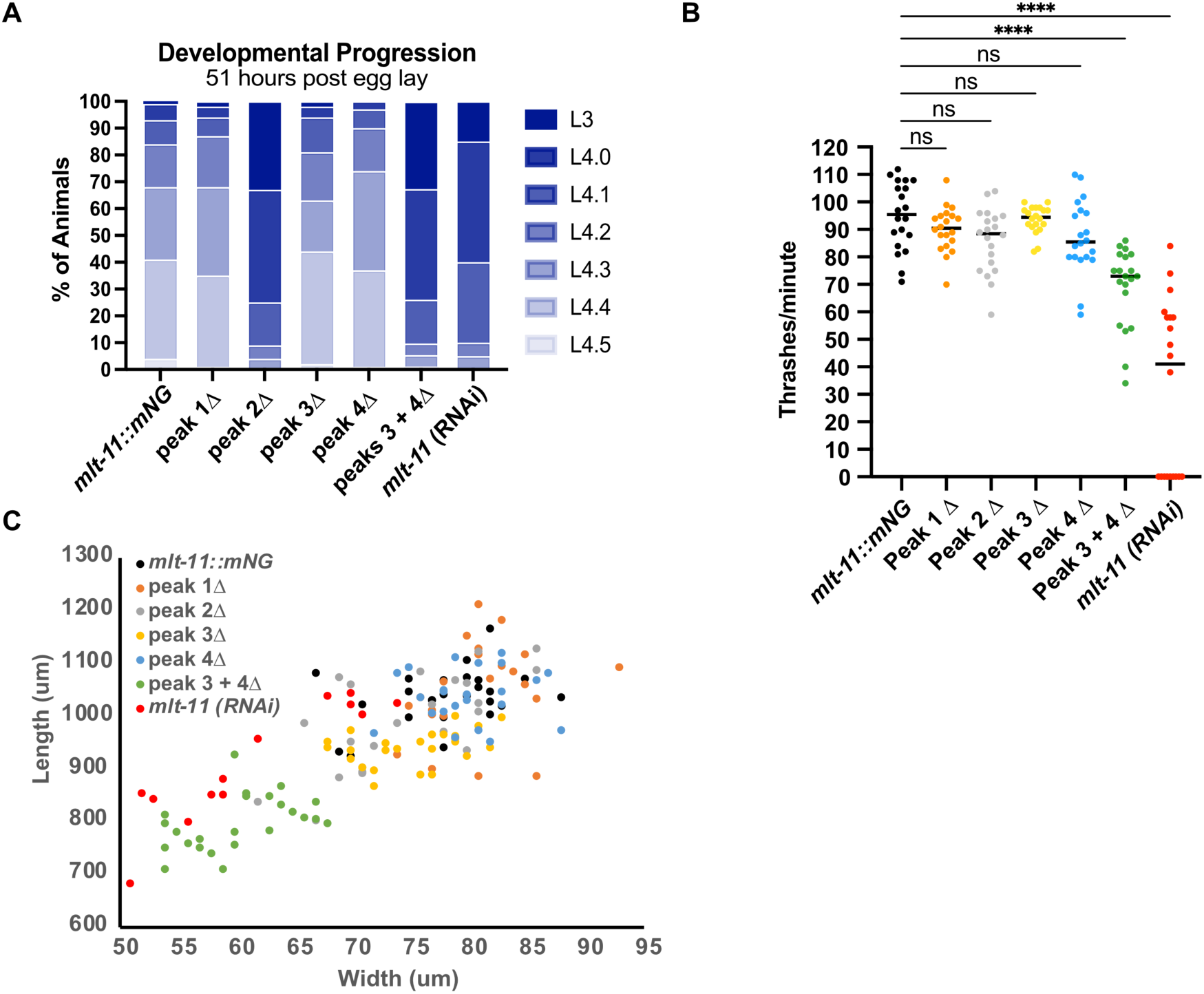
*mlt-11* expression abrogation causes developmental delay, motility defects and smaller bodies. Worms of the indicated genotypes and treatment (control or *mlt-11(RNAi))* were allowed to lay eggs for two hours, then removed from the plates. Embryos were allowed to develop for 51 hours, then scored for (A) developmental stage (L3 or L4 substage), (B) thrashes per minute and (C) body length and width.

### Reduction of *mlt-11* expression causes defective cuticle function and structure

Given that *mlt-11* expression is regulated by NHR-23 and that NHR-23 depletion causes a defective cuticle barrier, we next wished to see how *mlt-11* inactivation impacted cuticle structure and function. To test the cuticle barrier function, we incubated control, promoter deletion mutants and *mlt-11(RNAi)* animals with the cuticle impermeable, cell membrane permeable Hoechst 33258 dye and scored animals with stained nuclei. In wild-type control and single peak deletion animals we observed no Hoechst staining, while we observed staining in 91% of *peak 3+4Δ* deletion mutants and 93% of *mlt-11(RNAi)* worms (Fig. 5A). This barrier defect was comparable to the *bus-8* positive control strain (Partridge et al. 2008). Additionally, both *peak 3+4Δ* and *mlt-11(RNAi)* animals expressed an *nlp-29::GFP* promoter reporter generally activated by infection, acute stress, and physical damage to the cuticle (Fig. 5B) (Pujol et al. 2008; Zugasti and Ewbank 2009). Together, these indicate *mlt-11* is necessary for the barrier function of the cuticle.

**Fig. 5.**
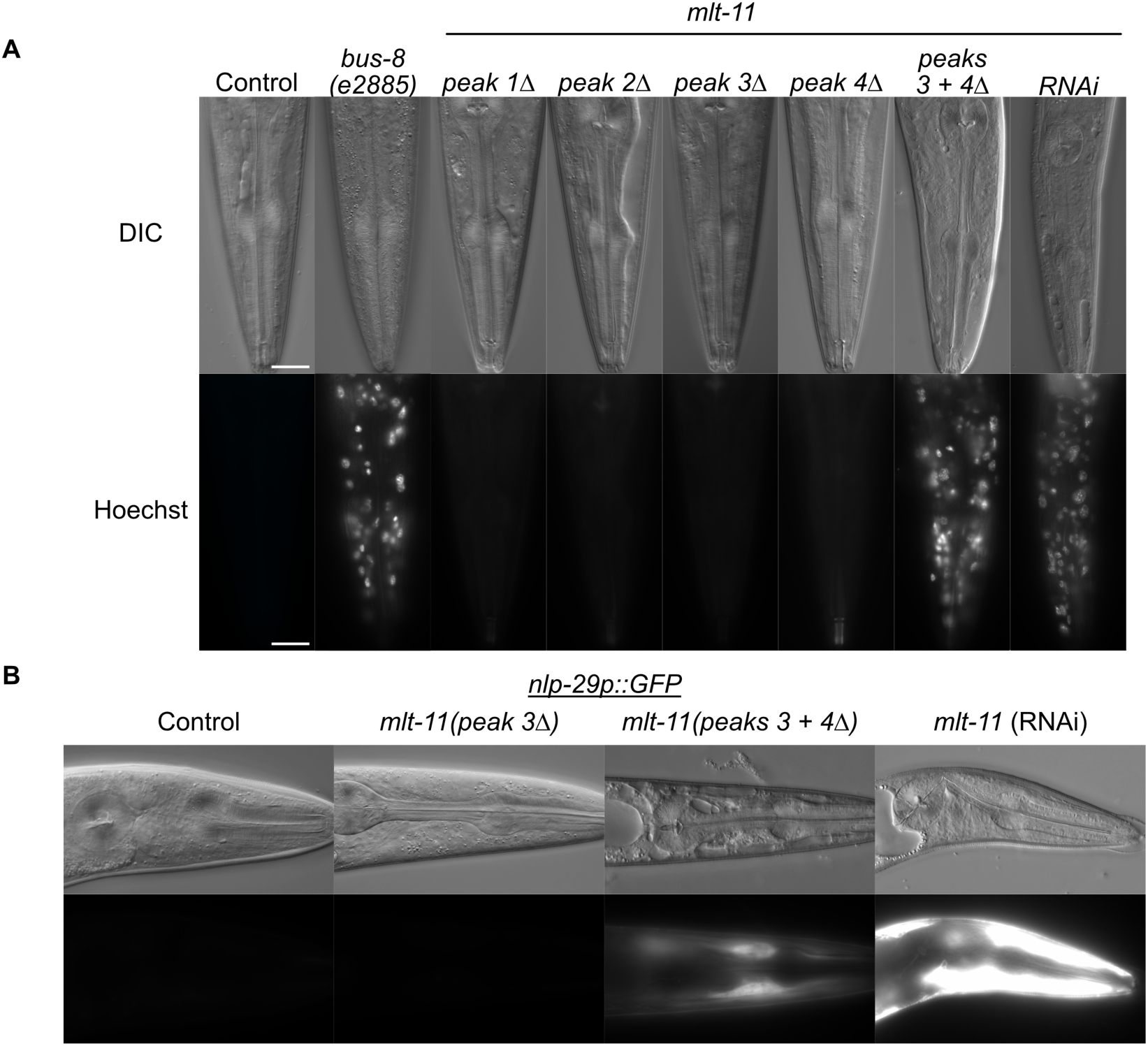
*mlt-11* knockdown causes defective aECM structure and function. (A,B) Representative images of mid-stage L4 larvae in animals of the indicated genotype/treatment. (A) Worms were washed and incubated with the cuticle impermeable/membrane permeable Hoechst 33258 dye before being imaged. (B) Worms carried an *nlp-29p::GFP* reporter activated by infection, acute stress, and physical damage to the cuticle [56,57]. Three biological replicates were performed and over 40 animals scored. Scale bars are 20 μm.

Given the barrier defect observed in *mlt-11(RNAi)* and *peak 3+4Δ* animals, we next wished to see how *mlt-11* inactivation impacted cuticle structure. The *C. elegans* cuticle is a complex structure that is composed of multiple layers (Fig. 6A)(Page and Johnstone 2007 Jan 1). In the adult cuticle, the basal layer is closest to the hypodermis, the cortical layer is furthest from the hypodermis and a fluid-filled medial layer is between them. Each layer contains a unique complement of collagen proteins that provide structure to the cuticle (Lažetić and Fay 2017). The components of the cuticle are secreted by hypodermal and seam cells and are assembled in distinct layers (G. N. Cox et al. 1981; Edgar et al. 1982; Page and Johnstone 2007 Jan 1). The cuticle secreted by the hypodermal syncytium form circumferential ridges called annuli separated from one another by furrows (Page and Johnstone 2007 Jan 1). In adults an inner basal layer contains two fibrous sub-layers angled in opposite directions and an outer cortical layer (Edgar et al. 1982). A fluid-filled medial layer contains hollow, nano-scale struts built from the BLI-1 and BLI-2 collagens (Edgar et al. 1982; Adams et al. 2023). All layers are composed of extensively cross-linked collagens. The cortical layer also contains cuticlins, proteins which remain in the insoluble fraction after cuticle solubilization (Ristoratore et al. 1994). The cortical layer is covered by the epicuticle, a poorly understood structure that is thought to be a lipid bilayer covered by a glycoprotein rich surface coat (Blaxter 1993; Peixoto and De Souza 1995; Bada Juarez et al. 2019). There can be stage-specific variations in cuticle structure. Only adults have a medial layer, and both L1 and adult cuticles contain alae, lateral longitudinal ridges secreted by seam cells in the epidermis (Edgar et al. 1982; Katz et al. 2022). Given that knockdown of *mlt-11* by RNAi (Frand *et al*., 2005) causes gross molting defects, we surmised that *mlt-11* could have a role in establishing or maintaining the structure of the cuticle, in apolysis or ecdysis. To test for a role in cuticle structure, we depleted *mlt-11* by RNAi or deleted promoter elements in strains carrying translational mNG fusions to proteins that mark the basal layer (ROL-6, COL-19), medial layer (BLI-1), and cortical layer (CUT-2) (Peixoto and De Souza 1995; Peixoto et al. 1998; Adams et al. 2023).

**Fig. 6.**
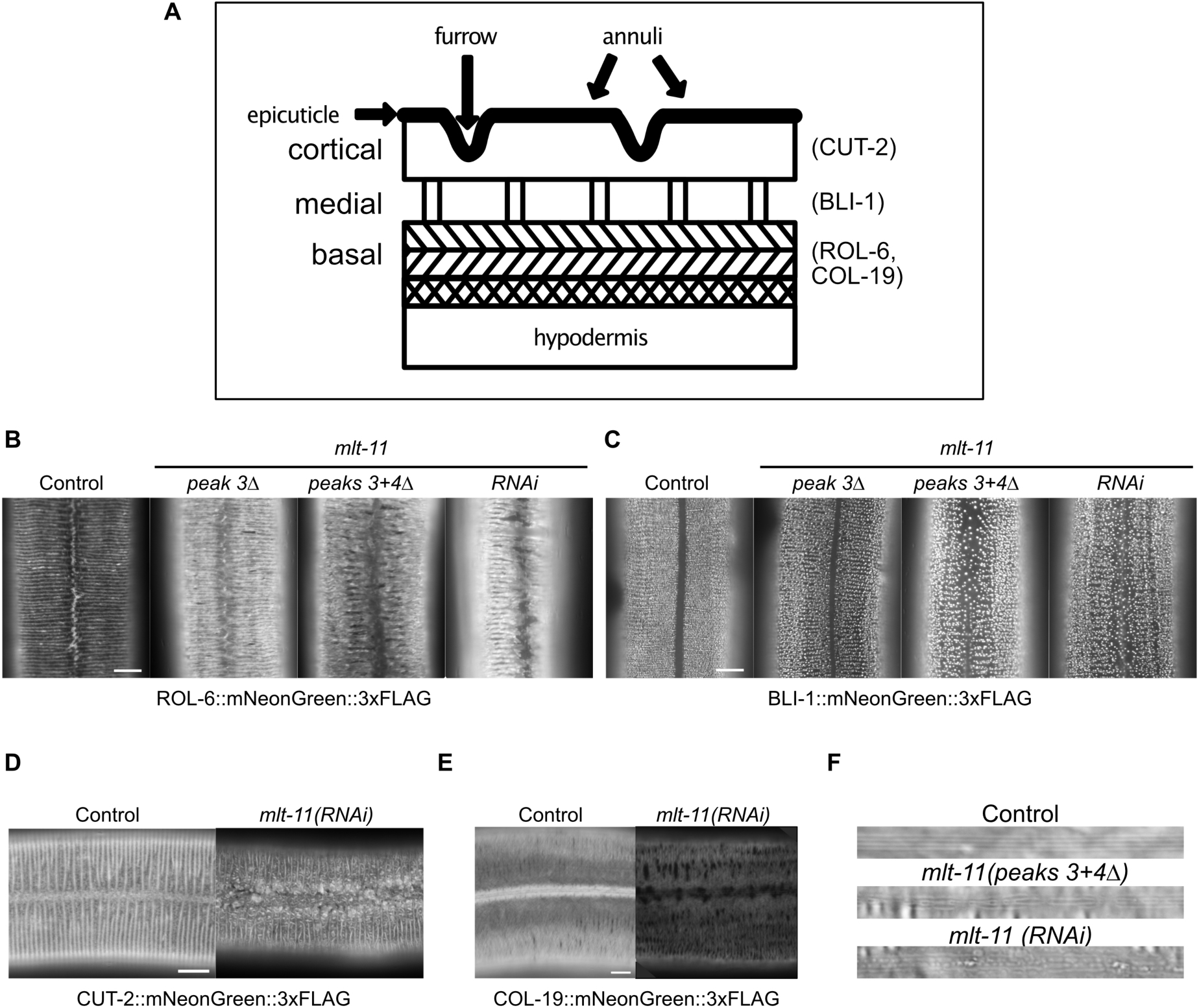
Disruption of *mlt-11* expression by RNAi or promoter deletion causes abnormalities in cuticle structure. (A) Cartoon model of the *C. elegans* adult cuticle sublayers (epicuticle, cortical, medial and basal), topological features (annuli and furrows). The localization of proteins used to mark specific layers of the cuticle in this study is indicated (CUT-2, BLI-1, ROL-6 and COL-19)[7,8,49]. Representative images of mNG::3xFLAG fused to ROL-6 (B), BLI-1 (C), CUT-2 (D) or COL-19 (E) in the cuticles of mid-stage L4 larvae in control (full promoter) or mlt-11 expression abrogated (peak 3Δ, peaks 3+4Δ, *mlt-11(RNAi)*) backgrounds. (F) (DIC images of adult alae in wild-type animals, *mlt-11(peaks 3+4Δ)* deletion mutants, and *mlt-11(RNAi)* animals. Scale bars are 10 μm in B-E and 5 μm in F.

ROL-6 is a collagen that localizes to tight longitudinal bands (furrows) in the basal layer of the adult cuticle (Fig. 6B)(Kim et al. 2010; Johnson et al. 2023). It also resides in patches along the junction of opposing annuli above seam cells (Fig. 6B). Following deletion of peak 3 or peaks 3+4, ROL-6::mNG localized to both furrows and annuli. Gaps were observed between annuli and patches were still seen, but smaller and fewer in number over seam cells. Following RNAi knockdown of *mlt-11*, annuli were separated by larger and more numerous gaps. Localization of ROL-6::mNG was absent over seam cells and opposing annuli at the junction were completely separated from each other and often broken at their termini (Fig. 6B). COL-19::mNG, another basal layer marker and adult-specific collagen normally localizes to alae and annuli, but following *mlt-11* RNAi we observed a reduction in expression and a loss of localization over the seam cells (Fig. 6E)(Thein et al. 2003).

BLI-1 is a structural collagen found in the medial layer of the adult cuticle, providing connection between the basal and cortical layers in the form of vertical struts (Tong et al. 2009; Adams et al. 2023). BLI-1::mNG localized in punctae organized in circumferential rows across the hyp7 cuticle, but is absent over seam cells (Fig. 6C). Deletion of *mlt-11* peak 3 led to gaps between the rows and inconsistency in the size of punctae (Fig. 6C). Deletion of peaks 3 and 4 together or RNAi knockdown caused further disorganization of BLI-1::mNG rows and mislocalization of punctae in seam cuticle (Fig. 6C). Similarly, CUT-2, a cuticlin in the cortical layer (Lassandro et al. 1994; Ristoratore et al. 1994), had localization patterns that were altered following knockdown of *mlt-11*. CUT-2::mNG displayed aberrant localization over the seam cells and a fibrous pattern over seam cells following *mlt-11* RNAi (Fig. 6D). Together, this suggests *mlt-11* is necessary for proper formation or patterning of each layer of the adult cuticle.

Given the aberrant localization of ROL-6, BLI-1, CUT-2, and COL-19 over the seam cells, we next examined alae morphology. The lateral alae are cuticle ridges formed by the interaction between the actin cytoskeleton in epithelial cells and an extracellular provisional matrix (George N. Cox et al. 1981; Katz et al. 2022). Three continuous longitudinal ridges span the midline of lateral surfaces of adult worms (Fig. 6F). *mlt-11(peak 3+4Δ)* animals had discontinuous alae and *mlt-11* RNAi produced a more severe phenotype (Fig. 6F). Together, these data indicate that *mlt-11* is necessary for accurate development of cuticle structures derived from both hypodermal and seam cells.

## DISCUSSION

Understanding how oscillating gene expression is coordinated during *C. elegans* molting will shed light on how this poorly understood biological timer functions. We confirmed that *mlt-11* is regulated by the NHR-23 transcription factor. We identified *cis-*regulatory elements in the *mlt-11* promoter sufficient for hypodermal and seam cell expression. Deletion of specific individual *cis-*regulatory elements (peak 3, peak4) in the endogenous reduced MLT-11::mNG expression, though did not produce significant defects. However, a *mlt-11(peak 3+4Δ)* mutant had a further reduction in MLT-11::mNG levels as well as developmental delay and cuticle structure and function defects. *mlt-11* inactivation affects the localization of translational fusions marking the basal (ROL-6::mNG, COL-19), medial (BLI-1::mNG), and cortical layers (CUT-2) of the cuticle indicating a broad role for MLT-11 in patterning this aECM.

### Distinct *cis-*regulatory elements control *mlt-11* expression

In our previous study of NHR-23, we examined the number of average NHR-23 peaks in oscillatory genes that peaked in expression in different phases (Johnson et al. 2023). Genes that peaked in expression in the hour following the *nhr-23* mRNA peak had the highest number of NHR-23 peaks flanking and within the gene body (Johnson et al. 2023). The average number of NHR-23 peaks declined as the peak phase occurred further from the *nhr-23* peak in expression (Johnson et al. 2023). One explanation for this trend is that genes that peak in expression closer to *nhr-23* could be more responsive to NHR-23 levels, similar to *E. coli* amino acid biosynthesis in which genes earlier in the pathway are more responsive (Zaslaver et al. 2004). We explored the regulation of *mlt-11* as it had six NHR-23 peaks flanking and within the gene body, and was a poorly characterized putative protease inhibitor; *nhr-23-*regulated genes were enriched in protease and protease inhibitor genes (Johnson et al. 2023). Using a promoter reporter-based strategy, we focused on the NHR-23 occupied regions upstream of the transcriptional start site. Peaks 3 and 4 drove robust reporter expression and NHR-23 controls the expression of a *mlt-11 peak 3* reporter (Fig. 2). In contrast, peak 1 drove undetectable expression and peak 2 drove weak expression primarily in seam cells (Fig. 2). These data are consistent with the heights of the four NHR-23 peaks (Fig. 2). While it is not clear whether NHR-23 regulates expression of these reporters through direct DNA binding or whether the regulation is indirect, there are consensus sites that NHR-23 is known to bind within or adjacent to these peak sequences (Fig. 2) (Kostrouchova et al. 1998). Consistent with the reporter data, deletion of the endogenous peak sequences revealed that peaks 3 and 4 produced the strongest effect on MLT-11::mNG expression and a double deletion of peaks 3 and 4 results in even stronger reduction in expression, suggesting an additive effect. Together, these data highlight that careful analysis of individual *cis-*regulatory elements is critical to dissect how NHR-23 controls the expression of its target genes. In a previous study, we showed that NOAH-1 localization but not level was altered by NHR-23 depletion despite having 8 NHR-23 peaks flanking and within its gene body (Johnson et al. 2023). The presence of an NHR-23 peak does not always indicate robust regulation.

### *mlt-11* is required for patterning of multiple cuticle layers

Mutation or depletion of oscillating collagens has been shown to affect the structural organization of collagens with similar temporal expression dynamics, suggesting these collagens might be part of a common substructure (McMahon et al. 2003). Surprisingly, *mlt-11* inactivation affected the localization of proteins in the basal, medial, and cortical layer of the cuticle (Fig. 6). Temporally, the *mlt-11* mRNA peaks in expression close to the *bli-1* expression peak and before the *rol-6* and *cut-2* mRNA peak (Meeuse et al. 2020). Thus, *mlt-11* affects the localization of proteins in multiple cuticle layers. MLT-11 has 10 predicted protease inhibitor domains most of which contain residues necessary for Kunitz inhibitor activity (Ascenzi et al. 2003; Ranasinghe and McManus 2013), which could buffer reduction of levels. It is possible that *mlt-11* mutant and knockdown phenotypes result from aberrant protease activity. Some proteases, such as BLI-4 and DPY-31 are thought to be involved in collagen processing in the secretory pathway, while others promote apolysis (Thacker et al. 1995; Davis et al. 2004; Stepek et al. 2010; Stepek et al. 2011; Birnbaum et al. 2023). Screening for genetic and protein-protein interactions between *mlt-11* and proteases is a high-priority future direction to determine how MLT-11 ensures proper cuticle structure/function.

Interestingly, we observed aberrant expression of ROL-6::mNG, CUT-2::mNG, and COL-19::mNG over the seam cells and alae defects (Fig. 6). Moreover, BLI-1 is normally excluded from the aECM over the seam cells but invades this area following *mlt-11* knockdown (Fig. 6). These data were reminiscent of our work on NHR-23 as depletion of this factor causes aberrant aECM formation over the seam cells (Johnson et al. 2023). Tissue-specific RNAi suggested that *nhr-23* activity also was necessary in the seam cells for developmental progression (Johnson et al. 2023). Exploring the roles of seam and hypodermal cells in constructing the cuticle is a future direction that we are focusing on.

### A *mlt-11* allelic series

Our RNAi and *cis-*regulatory element deletion experiments provided insight into the relationship between MLT-11 levels and phenotype. Following *cis-*regulatory element deletion in the *mlt-11* promoter and RNAi knockdown we observed a range of phenotypic severity: i) molting defects and developmental delay; iii) cuticle organization defects; and iii) cuticle barrier defects. These mutants and *mlt-11(RNAi)* comprise an allelic series to link *mlt-11* levels to specific phenotypes. Motility was also less sensitive to MLT-11 levels as we needed RNAi depletion to observe phenotypes (Fig. 4). BLI-1 localization, developmental rate and the cuticle barrier displayed intermediate sensitivity to MLT-11 levels as peak 3+4 deletion caused defects, as did RNAi (Fig. 4,5,6). Finally ROL-6 organization seemed highly sensitive to MLT-11 levels, as peak 3 deletion caused a reduction of MLT-11 levels and aberrant ROL-6 localization especially over the seam cells (Fig. 6). One interesting exception to the relationship between *mlt-11* levels and phenotype was that the *peak 2Δ* mutants had a developmental delay similar to *mlt-11(RNAi)* animals, but were otherwise wild type. Given that this *cis-*element seemed to primarily drive expression in the seam cells, this allele could hint at tissue-specific functions for MLT-11.

### Future perspective

Understanding the relative contribution of *mlt-11 cis-*regulatory elements to expression is an entry point to understanding the larger question of how genes involved in the molting cycle are regulated to peak at specific times during a given larval stage. Addressing this question will require an understanding of the transcriptional regulators of the molt cycle and how they are interconnected as well as detailed analysis of how these *trans*-factors interact with specific *cis-*elements to ensure that a given gene peaks at the correct time.

It will also be important to perform careful structure-function analyses to determine the role of specific Kunitz domains and other sequences in MLT-11 patterning of the cuticle. Pre-printed work suggests that the null phenotype is embryonic lethality, which would indicate that very little MLT-11 is required for embryonic development (Ragle et al. 2022). Understanding how MLT-11 patterns the cuticle and where it functions is a key future question.

## MATERIALS AND METHODS

### Strains and culture

*C. elegans* were cultured as originally described (Brenner 1974), except worms were grown on MYOB media instead of NGM. MYOB agar was made as previously described (Church et al. 1995).

### Strains created by injection in the Ward Lab and used in this study

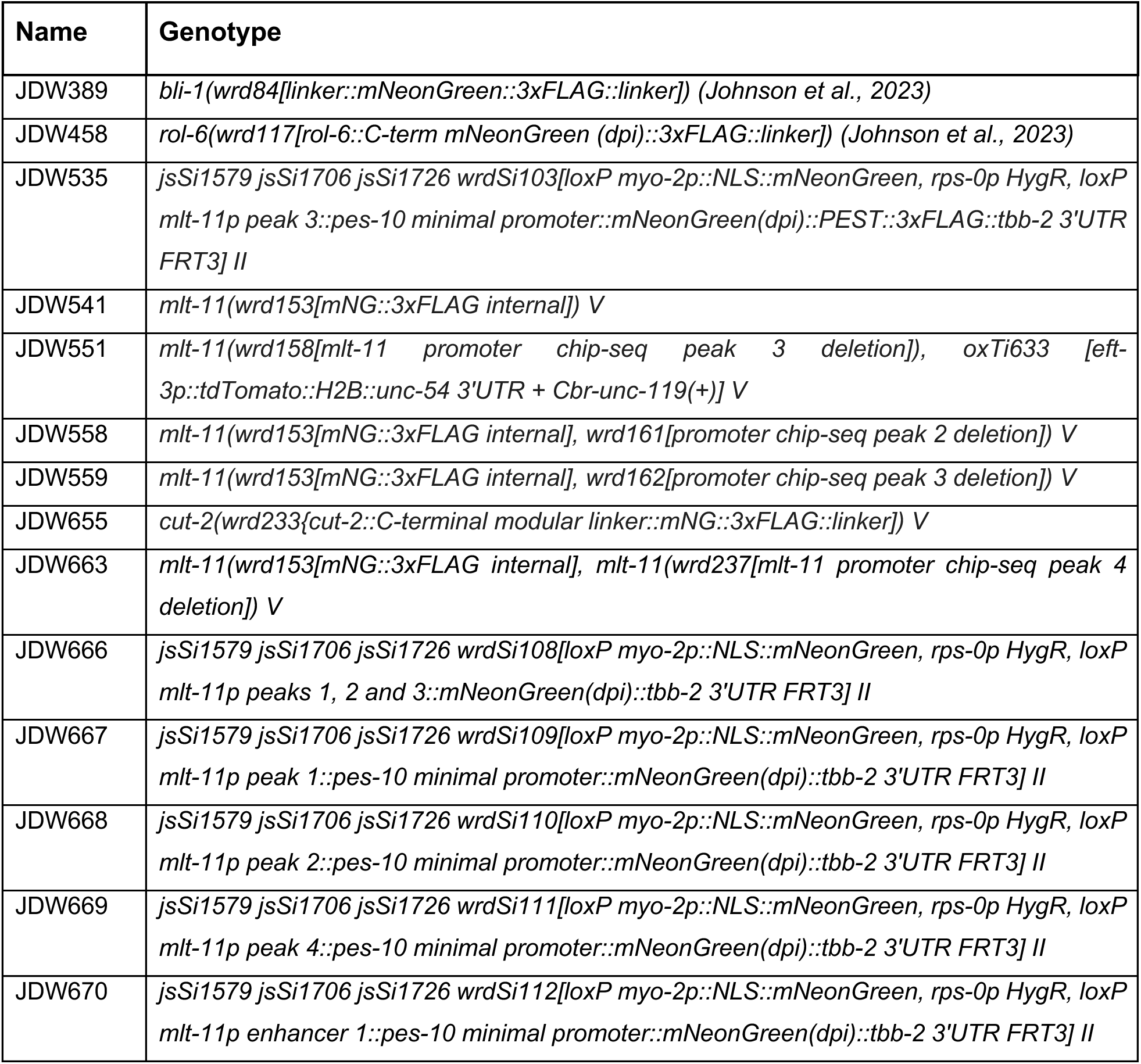
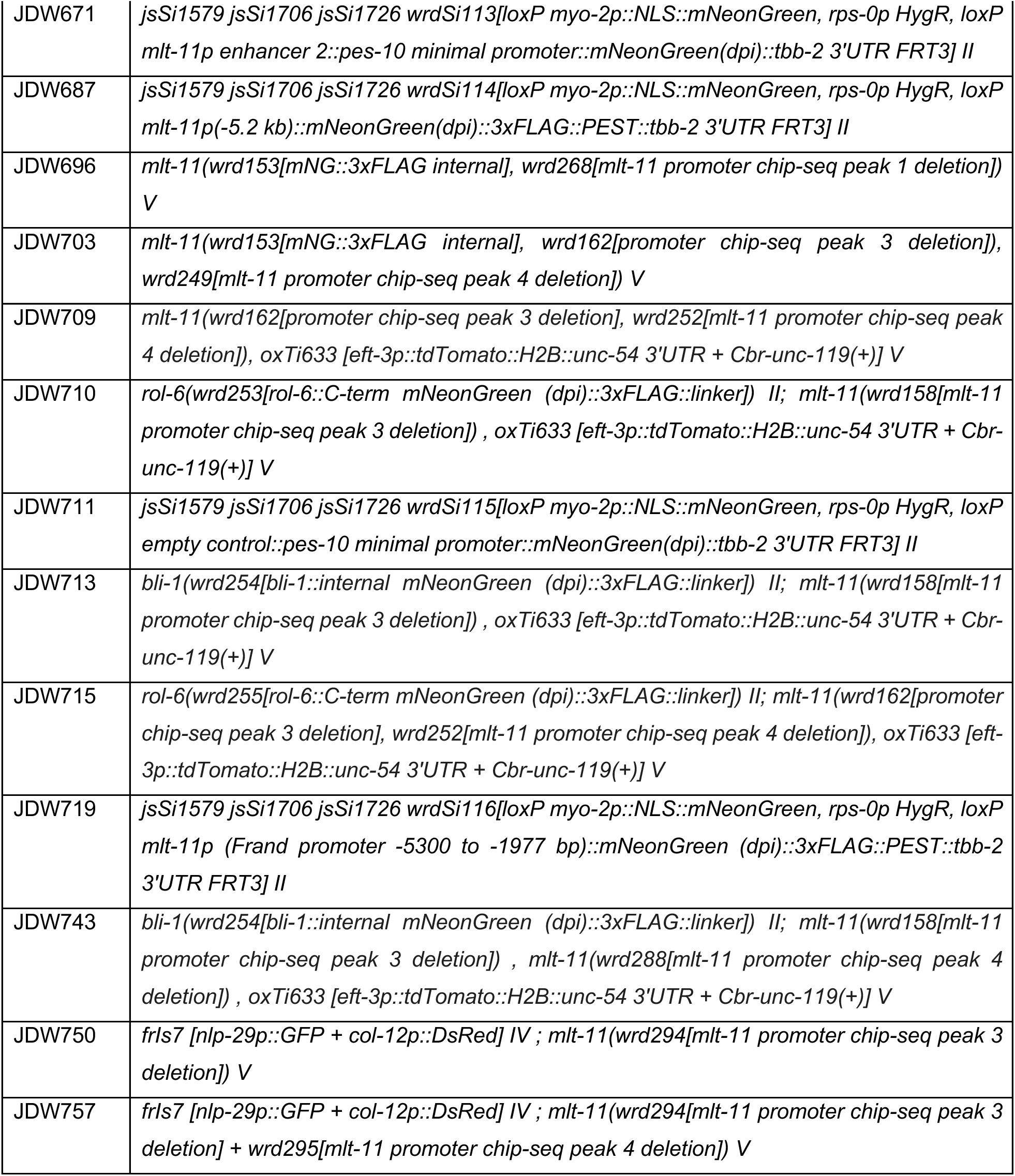

### Strains provided by the *Caenorhabditis* Genetics Center

Strains provided by the *Caenorhabditis* Genetics Center:
|  |  |
| --- | --- |
| N2 | Wild type |
| CB6193 | *bus-8B(e2885) X* (Partridge et al. 2008) |
| EG7968 | *unc-119(ed3) III; oxTi633 [eft-3p::tdTomato::H2B::unc-54 3'UTR + Cbr-unc-119(+)] V* (Frøkjær-Jensen et al. 2014) |
| IG274 | *frls7 [nlp-29p::GFP + col-12p::DsRed] IV* (Pujol et al. 2008) |
| NM5179 | *jsTi1493 [LoxP::mex-5p::FLP:SL2::mNeonGreen::rpl-28p::FRT::GFP::his-58::FRT3] IV* (Nonet 2020) |
| NM5548 | *jsSi1579 jsSi1706 jsSi1726[loxP myo-2p FRT nlsCyOFP myo-2 3' mex-5p FLP D5 glh-2 3' FRT3] II* (Nonet 2023) |

### Other strains

PHX4625 *col-19::mNG(cyb4625) X* was a gift from Dr. Andrew Chisholm and will be described elsewhere.

### Genome Editing

All plasmids used are listed in Table S1. Annotated plasmid sequence files are provided in File S1. Specific cloning details and primers used are available upon request. JDW380 *jsTi1493 {mosL loxP [wrdSi72(mlt-11(-2.8kb)p::mNeonGreen(dpi)::tbb-2 3’UTR)] FRT3::mosR} IV* was created by recombination-mediated cassette exchange (RMCE)(Nonet 2020). A 2.8 kb *mlt-11* promoter fragment was initially Gibson cloned into the *NLS::mScarlet (dpi)::tbb-2 3’UTR* vector pJW1841 (Ashley et al. 2021) to generate pJW1934. The mScarlet cassette was then replaced with mNeonGreen (dpi) to generate pJW2229. The *mlt-11p (−2.8kb) mNeonGreen (dpi)-tbb-2 3’UTR* fragment was PCR amplified from pJW2229 and Gibson cloned into *Sph*I-HF+*Spe*I-HF double digested RMCE integration vector pLF3FShC to produce pJW2337. This vector was integrated into NM5179 and the SEC was excised as previously described (Nonet 2020).

A pJW2361 *mNeonGreen(dpi)::3xFLAG::PEST-tbb-2 3’UTR* vector for SapTrap with ATG and GTA connectors was constructed by linearizing pJW2322 (Clancy et al. 2023 Feb 7) by PCR and Gibson cloning in PCR amplified *mNeonGreen (dpi)::3xFLAG* from pJW2172 and a *linker::PEST::tbb-2 3’UTR* from pJW1836 (Ashley et al. 2021). The pJW2286 *mlt-11p (−2.8 kb)* promoter for SapTrap cloning was previously described (Clancy et al. 2023 Feb 7). The *mlt-11* promoter fragment from Frand *et al*. (2005) and the *mlt-11p (−5.3 kb)* promoter fragment were PCR amplified from a fosmid containing the *mlt-11* gene and Gibson cloned into linearized pJW2286 to make pJW2451 and pJW2457, respectively. These *mlt-11* promoter plasmids were SapTrap cloned with pJW2361 and pNM4216 to generate insertion vectors for rapid RMCE (Schwartz and Jorgensen 2016; Nonet 2023). The remaining promoter reporters were constructed by SapTrap cloning and rapid RMCE (Schwartz and Jorgensen 2016; Nonet 2023). A pJW2365 *pes-10 minimal promoter::mNeonGreen(dpi):: 3xFLAG::PEST-tbb-2 3’UTR* vector for SapTrap with ATG and GTA connectors was constructed by linearing pJW2361 and Gibson cloning in a *pes-10* minimal promoter amplified from pJW1947 (Ashley et al. 2021). The ATAC-seq and NHR-23 ChIP-seq peak DNA sequences were PCR amplified from pJW2337 or pJW2457 and Gibson cloned into pDONR221 with TGG and ATG connectors for SapTrap. These plasmids were then SapTrap cloned with pJW2365 and pNM4216 to generate insertion vectors for rapid RMCE (Schwartz and Jorgensen 2016; Nonet 2023).

For the *mlt-11* internal knock-in (JDW541), the mNeonGreen::3xFLAG cassette was inserted in an unstructured region of exon 7. This strain was created by injection of RNPs [700 ng/µl IDT Cas9, 115 ng/µl crRNA and 250 ng/µl IDT tracrRNA] and a dsDNA repair template (25-50 ng/ul) created by PCR amplification of a pJW2172 plasmid template into N2 animals (Paix et al. 2014; Paix et al. 2015)(Table S1). PCR products were melted to boost editing efficiency, as previously described (Ghanta and Mello 2020). Sequences of CRISPR/Cas9-mediated genome edits are provided in File S2. crRNAs used are provided in Table S3 and repair template oligos for deletions are provided in Table S4. F1 progeny were screened by mNeonGreen expression.

*mlt-11* promoter region deletion strains were created by injection of Cas9 ribonucleoprotein complexes (RNPs)(Paix et al. 2014; Paix et al. 2015) [700 ng/µl IDT Cas9, 115 ng/µl each crRNA and 250 ng/µl IDT tracrRNA], oligonucleotide repair template (110 ng/µl) and pSEM229 co-injection marker (25 ng/µl)(El Mouridi et al. 2020) for screening into N2 or JDW541. Where possible, we selected “GGNGG” crRNA targets as these have been the most robust in our hand and support efficient editing (Farboud and Meyer 2015). F1s expressing the co-injection marker were isolated to lay eggs and screened by PCR for the deletion. Genotyping primers are provided in Table S2.

### Imaging

Synchronized animals were collected by either picking or washing off plates. For washing, 1000 µl of M9 + 2% gelatin was added to the plate or well, agitated to suspend animals in M9+gelatin, and then transferred to a 1.5 ml tube. Animals were spun at 700xg for 1 min. The media was then aspirated off and animals were resuspended in 500µl M9 + 2% gelatin with 5 mM levamisole. 12 µl of animals in M9 +gel with levamisole solution were placed on slides with a 2% agarose pad and secured with a coverslip. For picking, animals were transferred to a 10 µl drop of M9+5 mM levamisole on a 2% agarose pad on a slide and secured with a coverslip. Images were acquired using a Plan-Apochromat 40x/1.3 Oil DIC lens or a Plan-Apochromat 63x/1.4 Oil DIC lens on an AxioImager M2 microscope (Carl Zeiss Microscopy, LLC) equipped with a Colibri 7 LED light source and an Axiocam 506 mono camera. Acquired images were processed through Fiji software (version: 2.0.0-rc-69/1.52p). For direct comparisons within a figure, we set the exposure conditions to avoid pixel saturation of the brightest sample and kept equivalent exposure for imaging of the other samples.

### RNAi Knockdown

RNA interference experiments were performed as in Johnson *et al*. (2023). Control RNAi used an empty L4440. The *mlt-11 (RNAi)* vector was streaked from the Ahringer library (Kamath et al. 2003). Vector sequences are provided in File S1.

### Hoechst staining

Hoechst 33258 staining was performed as described previously (Moribe et al. 2004), except that we used 10 μg/ml of Hoechst 33258 as previously described (Ward et al. 2014). Two biological replicates were performed examining 50 animals per experiment. Representative images were taken with equivalent exposures using a 63× Oil DIC lens, as described in the imaging section.

### Data availability

A full description of all oligonucleotides, plasmids, transgenes, and *C. elegans* strains created and used in this article is in the set of supplemental tables. The authors affirm that all data necessary for confirming the conclusions of the article are present within the article, figures, tables, and Supplementary material. Plasmid sequences are provided in File S1, knock-in sequences from genome editing are provided in File S2. Any additional information is provided upon request.

## Acknowledgements

We would like to thank Prof. David Fay, Prof. Meera Sundaram, Prof. Andrew Chisholm, and John C. Clancy for helpful conversations. We thank Krista Myles, Patricia Bliatout, Zoe Johnson, Javier Hernandez Lopez, Zoie Reyna, Emma Cadena, Valarie Hallin, and Olivia Vedar for research support. We thank Prof. Andrew Chisholm and Prof. Michael Nonet for strains. Some strains were provided by the *Caenorhabditis* Genetics Center, which is funded by the NIH Office of Research Infrastructure Programs [P40 OD010440].

## Author Contributions

Conceptualization: J.M.R, J.D.W.

Methodology: J.M.R, A.T, A.J., A.A.V., V.P., J.D.W.

Validation: J.M.R, A.T, A.J., A.A.V., V.P., J.D.W.

Formal analysis: J.M.R, A.T, A.J., A.A.V., V.P., J.D.W.

Resources: J.M.R, J.D.W.

Data curation: J.M.R, J.D.W.

Writing - original draft: J.M.R, J.D.W.

Writing - review & editing: J.M.R, A.T, A.J., A.A.V., V.P., J.D.W.

Supervision: J.M.R, J.D.W.

Project administration: J.D.W.

Funding acquisition: J.D.W.

## Funding

This work was funded by the National Institutes of Health (NIH) National Institute

of General Medical Sciences (NIGMS) [R00GM107345 and R01GM138701] to J.D.W.

## REFERENCES

Adams JRG, Pooranachithra M, Jyo EM, Zheng SL, Goncharov A, Crew JR, Kramer JM, Jin Y, Ernst AM, Chisholm AD. 2023. Nanoscale patterning of collagens in *C. elegans* apical extracellular matrix. Nat Commun. 14(1):7506. doi:10.1038/s41467-023-43058-9.

Ascenzi P, Bocedi A, Bolognesi M, Spallarossa A, Coletta M, Cristofaro R, Menegatti E. 2003. The Bovine Basic Pancreatic Trypsin Inhibitor (Kunitz Inhibitor): A Milestone Protein. Curr Protein Pept Sci. 4(3):231–251. doi:10.2174/1389203033487180.

Ashley GE, Duong T, Levenson MT, Martinez MAQ, Johnson LC, Hibshman JD, Saeger HN, Palmisano NJ, Doonan R, Martinez-Mendez R, et al. 2021. An expanded auxin-inducible degron toolkit for *Caenorhabditis elegans*. Genetics. 217(3):iyab006. doi:10.1093/genetics/iyab006.

Bada Juarez J, O’Rourke D, Judge P, Liu L, Hodgkin J, Watts A. 2019. Lipodisqs for eukaryote lipidomics with retention of viability: Sensitivity and resistance to Leucobacter infection linked to C.elegans cuticle composition. Chem Phys Lipids. 222(2019).

Birnbaum SK, Cohen JD, Belfi A, Murray JI, Adams JRG, Chisholm AD, Sundaram MV. 2023. The proprotein convertase BLI-4 promotes collagen secretion prior to assembly of the *Caenorhabditis elegans* cuticle. PLOS Genet. 19(9):e1010944. doi:10.1371/journal.pgen.1010944.

Blaxter ML. 1993. Cuticle surface proteins of wild type and mutant *Caenorhabditis elegans*. J Biol Chem. 268(9):6600–6609.

Brenner S. 1974. The genetics of *Caenorhabditis elegans*. Genetics. 77(1):71–94.

Charlton WL, Harel HYM, Bakhetia M, Hibbard JK, Atkinson HJ, McPherson MJ. 2010. Additive effects of plant expressed double-stranded RNAs on root-knot nematode development. Int J Parasitol. 40(7):855–864. doi:10.1016/j.ijpara.2010.01.003.

Church DL, Guan KL, Lambie EJ. 1995. Three genes of the MAP kinase cascade, *mek-2*, *mpk-1/sur-1* and *let-60 ras*, are required for meiotic cell cycle progression in *Caenorhabditis elegans*. Development. 121(8):2525–2535.

Clancy JC, Vo AA, Myles KM, Levenson MT, Ragle JM, Ward JD. 2023 Feb 7. Experimental considerations for study of *C. elegans* lysosomal proteins. G3 GenesGenomesGenetics.:jkad032. doi:10.1093/g3journal/jkad032.

Cohen JD, Sundaram MV. 2020. *C. elegans* Apical Extracellular Matrices Shape Epithelia. J Dev Biol. 8(4):23. doi:10.3390/jdb8040023.

Cox G. N., Kusch M, Edgar RS. 1981. Cuticle of *Caenorhabditis elegans*: its isolation and partial characterization. J Cell Biol. 90(1):7–17. doi:10.1083/jcb.90.1.7.

Cox George N., Staprans S, Edgar RS. 1981. The cuticle of *Caenorhabditis elegans*: II. Stage-specific changes in ultrastructure and protein composition during postembryonic development. Dev Biol. 86(2):456–470. doi:10.1016/0012-1606(81)90204-9.

Davis MW, Birnie AJ, Chan AC, Page AP, Jorgensen EM. 2004. A conserved metalloprotease mediates ecdysis in *Caenorhabditis elegans*. Development. 131(23):6001–6008. doi:10.1242/dev.01454.

Edgar RS, Cox GN, Kusch M, Politz JC. 1982. The Cuticle of *Caenorhabditis elegans*. J Nematol. 14(2):248–258.

El Mouridi S, Peng Y, Frøkjær-Jensen C. 2020. Characterizing a strong pan-muscular promoter (P*mlc-1*) as a fluorescent co-injection marker to select for single-copy insertions. MicroPublication Biol. 2020(09). doi:10.17912/micropub.biology.000302.

Farboud B, Meyer BJ. 2015. Dramatic Enhancement of Genome Editing by CRISPR/Cas9 Through Improved Guide RNA Design. Genetics. 199(4):959–971. doi:10.1534/genetics.115.175166.

Frand AR, Russel S, Ruvkun G. 2005. Functional Genomic Analysis of *C. elegans* Molting. PLoS Biol. 3(10):e312. doi:10.1371/journal.pbio.0030312.

Frøkjær-Jensen C, Davis MW, Sarov M, Taylor J, Flibotte S, LaBella M, Pozniakovsky A, Moerman DG, Jorgensen EM. 2014. Random and targeted transgene insertion in *Caenorhabditis elegans* using a modified *Mos1* transposon. Nat Methods. 11(5):529–534. doi:10.1038/nmeth.2889.

Gerstein MB, Lu ZJ, Van Nostrand EL, Cheng C, Arshinoff BI, Liu T, Yip KY, Robilotto R, Rechtsteiner A, Ikegami K, et al. 2010. Integrative Analysis of the *Caenorhabditis elegans* Genome by the modENCODE Project. Science. 330(6012):1775–1787. doi:10.1126/science.1196914.

Ghanta KS, Mello CC. 2020. Melting dsDNA Donor Molecules Greatly Improves Precision Genome Editing in *Caenorhabditis elegans*. Genetics. 216(3):643–650. doi:10.1534/genetics.120.303564.

Gissendanner CR, Crossgrove K, Kraus KA, Maina CV, Sluder AE. 2004. Expression and function of conserved nuclear receptor genes in *Caenorhabditis elegans*. Dev Biol. 266(2):399–416. doi:10.1016/j.ydbio.2003.10.014.

Gissendanner CR, Sluder AE. 2000. *nhr-25*, the *Caenorhabditis elegans* ortholog of *ftz-f1*, is required for epidermal and somatic gonad development. Dev Biol. 221(1):259–272. doi:10.1006/dbio.2000.9679.

Hauser YP, Meeuse MWM, Gaidatzis D, Großhans H. 2021 Jul 5. The BLMP-1 transcription factor promotes oscillatory gene expression to achieve timely molting. bioRxiv.:2021.07.05.450828. 10.1101/2021.07.05.450828.

Hendriks G-J, Gaidatzis D, Aeschimann F, Großhans H. 2014. Extensive oscillatory gene expression during *C. elegans* larval development. Mol Cell. 53(3):380–392. doi:10.1016/j.molcel.2013.12.013.

Johnson LC, Vo AA, Clancy JC, Myles KM, Pooranachithra M, Aguilera J, Levenson MT, Wohlenberg C, Rechtsteiner A, Ragle JM, et al. 2023. NHR-23 activity is necessary for *C. elegans* developmental progression and apical extracellular matrix structure and function. Development. 150(10):dev201085. doi:10.1242/dev.201085.

Kamath RS, Fraser AG, Dong Y, Poulin G, Durbin R, Gotta M, Kanapin A, Le Bot N, Moreno S, Sohrmann M, et al. 2003. Systematic functional analysis of the *Caenorhabditis elegans* genome using RNAi. Nature. 421(6920):231–237. doi:10.1038/nature01278.

Katz SS, Barker TJ, Maul-Newby HM, Sparacio AP, Nguyen KCQ, Maybrun CL, Belfi A, Cohen JD, Hall DH, Sundaram MV, et al. 2022. A transient apical extracellular matrix relays cytoskeletal patterns to shape permanent acellular ridges on the surface of adult *C. elegans*. PLOS Genet. 18(8):e1010348. doi:10.1371/journal.pgen.1010348.

Kim D hyun, Grün D, van Oudenaarden A. 2013. Dampening of expression oscillations by synchronous regulation of a microRNA and its target. Nat Genet. 45(11):1337–1344. doi:10.1038/ng.2763.

Kim TH, Kim DH, Nam HW, Park SY, Shim J, Cho JW. 2010. Tyrosylprotein sulfotransferase regulates collagen secretion in *Caenorhabditis elegans*. Mol Cells. 29(4):413–418. doi:10.1007/s10059-010-0049-4.

Kostrouchova M, Krause M, Kostrouch Z, Rall JE. 1998. CHR3: a *Caenorhabditis elegans* orphan nuclear hormone receptor required for proper epidermal development and molting. Development. 125(9):1617–1626.

Kostrouchova M, Krause M, Kostrouch Z, Rall JE. 2001. Nuclear hormone receptor CHR3 is a critical regulator of all four larval molts of the nematode *Caenorhabditis elegans*. Proc Natl Acad Sci U S A. 98(13):7360–7365. doi:10.1073/pnas.131171898.

Kumar S, Chaudhary K, Foster JM, Novelli JF, Zhang Y, Wang S, Spiro D, Ghedin E, Carlow CKS. 2007. Mining Predicted Essential Genes of *Brugia malayi* for Nematode Drug Targets. PLoS ONE. 2(11):e1189. doi:10.1371/journal.pone.0001189.t003.

Lassandro F, Sebastiano M, Zei F, Bazzicalupo P. 1994. The role of dityrosine formation in the crosslinking of CUT-2, the product of a second cuticlin gene of *Caenorhabditis elegans*. Mol Biochem Parasitol. 65(1):147–159. doi:10.1016/0166-6851(94)90123-6.

Lažetić V, Fay DS. 2017. Molting in *C. elegans*. Worm. 6(1):1–20. doi:10.1080/21624054.2017.1330246.

McMahon L, Muriel JM, Roberts B, Quinn M, Johnstone IL. 2003. Two sets of interacting collagens form functionally distinct substructures within a *Caenorhabditis elegans* extracellular matrix. Mol Biol Cell. 14(4):1366–1378. doi:10.1091/mbc.e02-08-0479.

Meeuse MW, Hauser YP, Morales Moya LJ, Hendriks G, Eglinger J, Bogaarts G, Tsiairis C, Großhans H. 2020. Developmental function and state transitions of a gene expression oscillator in *Caenorhabditis elegans*. Mol Syst Biol. 16(7):e9975. doi:10.15252/msb.20209498.

Meeuse MWM, Hauser YP, Nahar S, Smith AAT, Braun K, Azzi C, Rempfler M, Großhans H. 2023. *C. elegans* molting requires rhythmic accumulation of the Grainyhead/LSF transcription factor GRH-1. EMBO J. doi:10.15252/embj.2022111895.

Meng J, Ma X, Tao H, Jin X, Witvliet D, Mitchell J, Zhu M, Dong M-Q, Zhen M, Jin Y, et al. 2017. Myrf ER-Bound Transcription Factors Drive *C. elegans* Synaptic Plasticity via Cleavage-Dependent Nuclear Translocation. Dev Cell. 41(2):180–194.e7. doi:10.1016/j.devcel.2017.03.022.

Miao R, Li M, Zhang Q, Yang C, Wang X. 2020. An ECM-to-Nucleus Signaling Pathway Activates Lysosomes for *C. elegans* Larval Development. Dev Cell. 52(1):21–37.e5. doi:10.1016/j.devcel.2019.10.020.

Moribe H, Yochem J, Yamada H, Tabuse Y, Fujimoto T, Mekada E. 2004. Tetraspanin protein (TSP-15) is required for epidermal integrity in *Caenorhabditis elegans*. J Cell Sci. 117:5209–5220. doi:10.1242/jcs.01403.

Nonet ML. 2020. Efficient Transgenesis in *Caenorhabditis elegans* Using Flp Recombinase-Mediated Cassette Exchange. Genetics. 215(4):903–921. doi:10.1534/genetics.120.303388.

Nonet ML. 2023. Rapid generation of *Caenorhabditis elegans* single-copy transgenes combining recombination-mediated cassette exchange and drug selection. Genetics. 224(3):iyad072. doi:10.1093/genetics/iyad072.

Page AP, Johnstone IL. 2007 Jan 1. The cuticle. WormBook Online Rev C. elegans Biol.:1–15. doi:10.1895/wormbook.1.138.1.

Page AP, Stepek G, Winter AD, Pertab D. 2014. Enzymology of the nematode cuticle: A potential drug target? Int J Parasitol DRUGS DRUG Resist. 4(2):133–141. doi:10.1016/j.ijpddr.2014.05.003.

Paix A, Folkmann A, Rasoloson D, Seydoux G. 2015. High Efficiency, Homology-Directed Genome Editing in *Caenorhabditis elegans* Using CRISPR-Cas9 Ribonucleoprotein Complexes. Genetics. 201(1):47–54. doi:10.1534/genetics.115.179382.

Paix A, Wang Y, Smith HE, Lee C-YS, Calidas D, Lu T, Smith J, Schmidt H, Krause MW, Seydoux G. 2014. Scalable and versatile genome editing using linear DNAs with microhomology to Cas9 Sites in *Caenorhabditis elegans*. Genetics. 198(4):1347–1356. doi:10.1534/genetics.114.170423.

Partridge FA, Tearle AW, Gravato-Nobre MJ, Schafer WR, Hodgkin J. 2008. The *C. elegans* glycosyltransferase BUS-8 has two distinct and essential roles in epidermal morphogenesis. Dev Biol. 317(2):549–559. doi:10.1016/j.ydbio.2008.02.060.

Peixoto CA, De Souza W. 1995. Freeze-fracture and deep-etched view of the cuticle of *Caenorhabditis elegans*. Tissue Cell. 27(5):561–568. doi:10.1016/s0040-8166(05)80065-5.

Peixoto CA, de Melo JV, Kramer JM, de Souza W. 1998. Ultrastructural Analyses of the *Caenorhabditis elegans rol-6 (su1006)* Mutant, Which Produces Abnormal Cuticle Collagen. J Parasitol. 84(1):45–49. doi:10.2307/3284528.

Pujol N, Cypowyj S, Ziegler K, Millet A, Astrain A, Goncharov A, Jin Y, Chisholm AD, Ewbank JJ. 2008. Distinct innate immune responses to infection and wounding in the *C. elegans* epidermis. Curr Biol. 18(7):481–489. doi:10.1016/j.cub.2008.02.079.

Ragle JM, Aita AL, Morrison KN, Martinez-Mendez R, Saeger HN, Ashley GA, Johnson LC, Schubert KA, Shakes DC, Ward JD. 2020. The conserved molting/circadian rhythm regulator NHR-23/NR1F1 serves as an essential co-regulator of *C. elegans* spermatogenesis. Development. 147(22):dev193862. doi:10.1242/dev.193862.

Ragle JM, Levenson MT, Clancy JC, Vo AA, Pham V, Ward JD. 2022. The conserved, secreted protease inhibitor MLT-11 is necessary for *C. elegans* molting and embryogenesis.:2022.06.29.498124. doi:10.1101/2022.06.29.498124.

Ragle JM, Morrison KN, Vo AA, Johnson ZE, Hernandez Lopez J, Rechtsteiner A, Shakes DC, Ward JD. 2022 Sep 22. NHR-23 and SPE-44 regulate distinct sets of genes during *C. elegans* spermatogenesis. G3 Bethesda Md.:jkac256. doi:10.1093/g3journal/jkac256.

Ranasinghe S, McManus DP. 2013. Structure and function of invertebrate Kunitz serine protease inhibitors. Dev Comp Immunol. 39(3):219–227. doi:10.1016/j.dci.2012.10.005.

Ristoratore F, Cermola M, Nola M, Bazzicalupo P, Favre R. 1994. Ultrastructural immuno-localization of CUT-1 and CUT-2 antigenic sites in the cuticles of the nematode *Caenorhabditis elegans*. J Submicrosc Cytol Pathol. 26(3):437–443.

Russel S, Frand AR, Ruvkun G. 2011. Regulation of the *C. elegans* molt by *pqn-47*. Dev Biol. 360(2):297–309. doi:10.1016/j.ydbio.2011.09.025.

Schwartz ML, Jorgensen EM. 2016. SapTrap, a Toolkit for High-Throughput CRISPR/Cas9 Gene Modification in *Caenorhabditis elegans*. Genetics. 202(4):1277–1288. doi:10.1534/genetics.115.184275.

Singh RN, Sulston JE. 1978. Some observations on molting in *C. elegans*. Nematologica. 24:63–71.

Stec N, Doerfel K, Hills-Muckey K, Ettorre VM, Ercan S, Keil W, Hammell CM. 2021. An Epigenetic Priming Mechanism Mediated by Nutrient Sensing Regulates Transcriptional Output during *C. elegans* Development. Curr Biol. 31(4):809–826.e6. doi:10.1016/j.cub.2020.11.060.

Stepek G, McCormack G, Birnie AJ, Page AP. 2011. The astacin metalloprotease moulting enzyme NAS-36 is required for normal cuticle ecdysis in free-living and parasitic nematodes. Parasitology. 138(2):237–248. doi:10.1017/S0031182010001113.

Stepek G, McCormack G, Page AP. 2010. Collagen processing and cuticle formation is catalysed by the astacin metalloprotease DPY-31 in free-living and parasitic nematodes. Int J Parasitol. 40(5):533–542. doi:10.1016/j.ijpara.2009.10.007.

Sundaram MV, Pujol N. 2024. The *Caenorhabditis elegans* cuticle and precuticle: a model for studying dynamic apical extracellular matrices in vivo. Genetics. 227(4):iyae072. doi:10.1093/genetics/iyae072.

Thacker C, Peters K, Srayko M, Rose AM. 1995. The *bli-4* locus of *Caenorhabditis elegans* encodes structurally distinct kex2/subtilisin-like endoproteases essential for early development and adult morphology. Genes Dev. 9(8):956–971. doi:10.1101/gad.9.8.956.

Thein MC, McCormack G, Winter AD, Johnstone IL, Shoemaker CB, Page AP. 2003. *Caenorhabditis elegans* exoskeleton collagen COL-19: an adult-specific marker for collagen modification and assembly, and the analysis of organismal morphology. Dev Dyn. 226(3):523–539. doi:10.1002/dvdy.10259.

Tong A, Lynn G, Ngo V, Wong D, Moseley SL, Ewbank JJ, Goncharov A, Wu Y-C, Pujol N, Chisholm AD. 2009. Negative regulation of *Caenorhabditis elegans* epidermal damage responses by death-associated protein kinase. Proc Natl Acad Sci U S A. 106(5):1457– 1461. doi:10.1073/pnas.0809339106.

Ward JD, Mullaney B, Schiller BJ, He LD, Petnic SE, Couillault C, Pujol N, Bernal TU, Van Gilst MR, Ashrafi K, et al. 2014. Defects in the *C. elegans* acyl-CoA Synthase, *acs-3*, and Nuclear Hormone Receptor, *nhr-25*, Cause Sensitivity to Distinct, but Overlapping Stresses. PLoS ONE. 9(3):e92552. doi:10.1371/journal.pone.0092552.s013.

Zaslaver A, Mayo AE, Rosenberg R, Bashkin P, Sberro H, Tsalyuk M, Surette MG, Alon U. 2004. Just-in-time transcription program in metabolic pathways. Nat Genet. 36(5):486–491. doi:10.1038/ng1348.

Zugasti O, Ewbank JJ. 2009. Neuroimmune regulation of antimicrobial peptide expression by a noncanonical TGF-β signaling pathway in *Caenorhabditis elegans* epidermis. Nat Immunol. 10(3):249–256. doi:10.1038/ni.1700.

